# Dynamics of SARS-CoV-2 genetic mutations and their information entropy

**DOI:** 10.1101/2022.06.13.495895

**Authors:** Melvin M. Vopson

## Abstract

We report an investigation of the mutations dynamics of the SARS-CoV-2 virus using Shannon’s information theory. Our study includes seventeen RNA genetic sequences collected at different geographic locations and timeframes ranging from Dec. 2019 to Oct. 2021. The data shows a previously unobserved relationship between the information entropy of genomes and their mutation dynamics. The information entropy of the mutated variants decreases linearly with the number of genetic mutations with a negative slope of 1.52 × 10^-5^ bits / mutations, pointing to a possible deterministic approach to the dynamics of genetic mutations. The method proposed here could be used to develop a predictive algorithm of genetic mutations.

---

The DNA or RNA genetic molecule is a highly complex information encoding system that displays incredible properties such as extremely low error rate, high stability, ability to self-replicate, self-repair, and to produce other molecules. The biological information is encoded in the sequence of nucleotides and the time evolution of a genome is described by the occasionally induced genetic mutations. Mutations are errors that appear during the process of self-replication when the genome is damaged and left unrepaired. These errors manifest in changes in the genetic sequence ranging from simple substitutions that maintain the number of nucleotides constant, to deletions and insertions that shift the sequence frame to shorter or larger size, respectively. Genetic mutations can affect an organism’s phenotype and are essential for the diversity and the evolution of living organisms. Because of the protective mechanisms like DNA / RNA self-repair or proof reading during replication, mutation rates are very low, but avoiding mutations could become too energetically costly to living organisms and they never reach zero. The scientific consensus on the dynamics of genetic mutations is the Darwinian theory of evolution, in which genetic mutations are random processes [1]. This means that mutations occur randomly regardless whether an organism will benefit or not from the DNA / RNA changes. Only the natural selection determines which mutations are beneficial and preserved in the course of evolution [2], but there is no observed correlation between any parameter or variable and the probability that these mutations will occur or not.

A recent study reported that mutations in the plant *Arabidopsis thaliana* occur less often in functionally constrained regions of the genome, challenging the paradigm that a directionless force in evolution drives mutations and proposing that an adaptive mutation bias exists [3].

Nei also concluded that the driving force of phenotypic evolution is mutation, and natural selection is of secondary importance [4].

While the existence of an adaptive mutation force is indeed a strong contra argument to the evolutionary theory that mutations occur randomly with respect to their consequences, one has no means of predicting future genetic mutations without prior knowledge of the exact adaptive path that an organism would follow, which is still an unknown outcome.

Using a mathematical – physical approach, here we report solid evidence of a truly predictive and quantifiable pattern of the dynamics of genetic mutations and we formulate a possible governing law of genetic mutations.

## Results

### Information theory applied to genomics

Our mathematical-physical analysis of genomic sequences is based on the Shannon’s information theory [5] and the concept of information entropy (IE). For a given set of n independent and distinctive events *Y = {y_1_, y_2_,…,y_m_}*, which are described by a discrete probability distribution *P = {p_1_, p_2_,…,p_m_}* on *Y*, the information content per event measured in bits, or the IE extracted when observing the set of events Y once is:

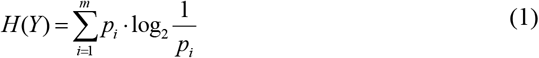

where the function *H(Y)* is called the Shannon IE and it is maximum when the events y_j_ have equal probabilities of occurring, *p_i_* =1/ *m*, so *H* (*Y*) =log_2_*m*. Observing N sets of events Y, or N times the set of events Y, the information extracted from the observation is N·H(Y) bits. A DNA sequence can be represented as a long string of the letters A, C, G, and T. These represent the four nucleotides: adenine (A), cytosine (C), guanine (G), and thymine (T) (replaced with uracil (U) in RNA sequences). Hence, within Shannon’s information theory framework, a typical genome can be represented as a 4-state probabilistic system, with *m = 4* distinctive events*, Y = {A, C, G,T*} and probabilities *p*= {*p_A_*, *p_C_*, *p_G_*, *p_T_*}. Assuming the events have equal probabilities of occurring, using digital information units, i.e. base of the logarithm is 2, and equation (1) for *m = 4*, we determine that maximum Shannon IE is 2, indicating that the maximum information encode per nucleotide is 2 bits, i.e. *A = 00, C = 01, G = 10, T = 11.*

Thinking of the genome as a non-equilibrium information coding system with a very long relaxation time, and given the highly complex nature of genome architecture, the main assumption here is that the IE profile of genetic sequences may represent a characteristic pattern able to highlight characteristic changes at sites where the nucleotide sequence is changed, such as single nucleotide polymorphisms (SNPs), deletions and insertions, suggesting not only a role for IE in the determination of genomic variation as already proven [6], but also a role in developing a deterministic approach to genetic mutations.

### IE of SARS-CoV-2 genomes

In order to test this assumption, we need to examine the time evolution of the IE of a genome that undergoes frequent mutations in a short period of time. A system that fits perfectly this requirement is a virus genome, and we examined the RNA sequence of the novel SARS-CoV-2 virus, which emerged in Dec. 2019 resulting in the current COVID-19 pandemic.

The reference RNA sequence of the SARS-CoV-2, representing a sample of the virus collected early in the pandemic in Wuhan, China in December 2019 (MN908947) [7], has 29,903 nucleotides, so N = 29,903. For this reference sequence we computed the Shannon IE using relation (1) and previously developed software, GENIES [8,9], designed to study the genetic mutations using Shannon’s IE [6]. The value obtained represents the reference Shannon IE at time zero, before any mutations took place. Using the NCBI virus database, we searched and extracted a number of SARS-CoV-2 variants sequenced at various locations around the globe, at different times, starting from Jan. 2020 to Oct. 2021. Our search was centred on complete genome sequences containing the same number of nucleotides as the reference sequence. We also selected carefully variants that displayed incremental number of SNP mutations, and we computed the Shannon IE corresponding to each variant. The full data set, including genome data references / links, collection times, number of SNP mutations, and the Shannon IE value of each genome are shown in Table 1. Figure 1 shows the computed IE of each SARS-CoV-2 variant as a function of its number of SNP mutations. Remarkably, the data indicates that the Shannon IE of the RNA genomes decreases linearly with their corresponding number of mutations. The larger is the number of mutations, the lower is the corresponding Shannon IE value. All analysed variants displayed the universal behaviour of lowering the IE value relative to that of the original genome.

**Figure 1.**
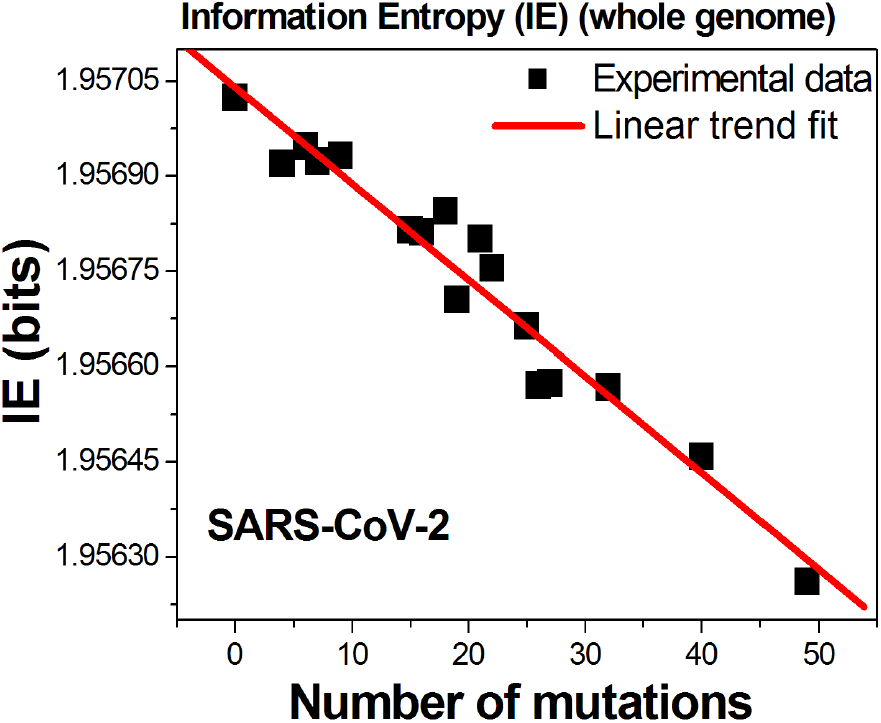
Information entropy values of variants of the SARS-CoV-2 virus as a function of the SNP mutations detected per variant.

**Table 1.** Tabulated results of the analysis performed on selected SARS-CoV-2 variants sequenced at various locations around the globe, over a period of 22 months.

| Genome | Ref. | SNPs | Collection time | Location | Shannon IE (bits) |
| --- | --- | --- | --- | --- | --- |
| MN908947 | [7] | 0 | Dec. 2019 | China | 1.9570243 |
| LC542809 | [10] | 4 | Mar. 2020 | Japan | 1.9569197 |
| MT956916 | [11] | 6 | May 2020 | Spain | 1.9569468 |
| MT956915 | [12] | 7 | May 2020 | Spain | 1.9569230 |
| MW466798 | [13] | 9 | Jul. 2020 | South Korea | 1.9569327 |
| OM098415 | [14] | 15 | May 2021 | China | 1.9568149 |
| OM098426 | [15] | 16 | May 2021 | China | 1.9568117 |
| MW679504 | [16] | 18 | Feb. 2021 | USA | 1.9568451 |
| MW294011 | [17] | 19 | Oct. 2020 | Ecuador | 1.9567058 |
| OK246811.1 | [18] | 21 | Mar. 2021 | USA / PA | 1.9568012 |
| OK246805 | [19] | 22 | Mar 2021 | USA / PA | 1.9567543 |
| MW679505 | [20] | 25 | Feb. 2021 | USA | 1.9566630 |
| MW735975 | [21] | 26 | Feb. 2021 | USA / OH | 1.9565714 |
| MW695383 | [22] | 27 | Dec. 2020 | USA / VA | 1.9865744 |
| OK546282.1 | [23] | 32 | Apr. 2021 | USA / NE | 1.9565675 |
| OK104651.1 | [24] | 40 | Aug. 2021 | Egypt | 1.9564591 |
| OL351371.1 | [25] | 49 | Oct. 2021 | Egypt | 1.9562614 |

The data in figure 1 has been selected to highlight this observation. Since the general consensus is that genetic mutations are random processes, one should not detect any correlation between the evolution of the genetic mutations and any other parameter.

The linear fit indicates that the link between the IE of genomes and the number of genetic mutations, n, is given by:

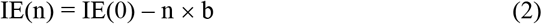

where IE(0) is the information entropy of the original genome before any mutations took place and b is the slope, b = 15.2 μbits / mutations for this particular study.

### The governing entropic law of genetic mutations

Our novel data indicates that there is an intimate link between the dynamics of the genetic mutations and the mathematical value of their IE measured in bits. This interesting result naturally stimulates the thought that perhaps there is a hidden and undiscovered deterministic approach to genetic mutations. In fact, the observed trend has a universal applicability to all information systems, beyond biological DNA / RNA information. It appears that all information systems are governed by an entopic law similar to the second law of thermodynamics, called the second law of infodynamics, which will be reported in a different article [26]. However, based on the data presented here, we are able to conclude this study by formulating a possible governing law of genetic mutation:

> *Biological RNA and DNA information systems tend to reduce their information entropy value over time by undergoing genetic mutations.*

### Predicting mutations from IE spectra

The careful application of this law facilitates a possible predictive approach to future genetic mutations, before they take place. This is a matter of future studies and beyond the scope of this report, but a possible way of taking advantage of the observed law could be a methodology that involves splitting the entire genome into subsets, called “windows”. This is because our analysis has been applied to the entire genetic sequences, each containing 29,903 characters, and the method gives no relationship between the IE value and the location of the mutations within the genome sequence, making the possibility of finding a full predictive algorithm of future genetic mutations unachievable yet. A “window” contains a given number of nucleotides (characters) called “window size” (WS). The windows representation of the genome is obtained by sliding the window from left to right, across the whole genome [6]. Each new position of the window is obtained by sliding it for a fixed number of characters, called the “step size” (SS). In order to ensure that all sections of the genome are captured by this process, the SS must be at least 1 and maximum WS, so 1< SS ≤WS. By doing this, a given genome of N nucleotides will result in a total of N_w_ windows, given by the formula:

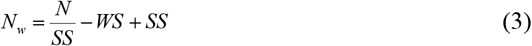

In order to produce an IE profile of the whole genome, one needs to compute the IE value of each window and to plot the IE values obtained as a function of the window index location within the genome. The result is an IE spectrum, which is a numerical representation of the genome sequence containing all its biological information converted into a numerical spectrum. Using GENIES [8,9] we computed the IE spectra (see examples in figure 3) by counting and working out the probabilities of occurrence of single nucleotides for SS = WS = 103. Figure 2 shows the average IE per window of each SARS-CoV-2 spectrum for three

**Figure 2.**
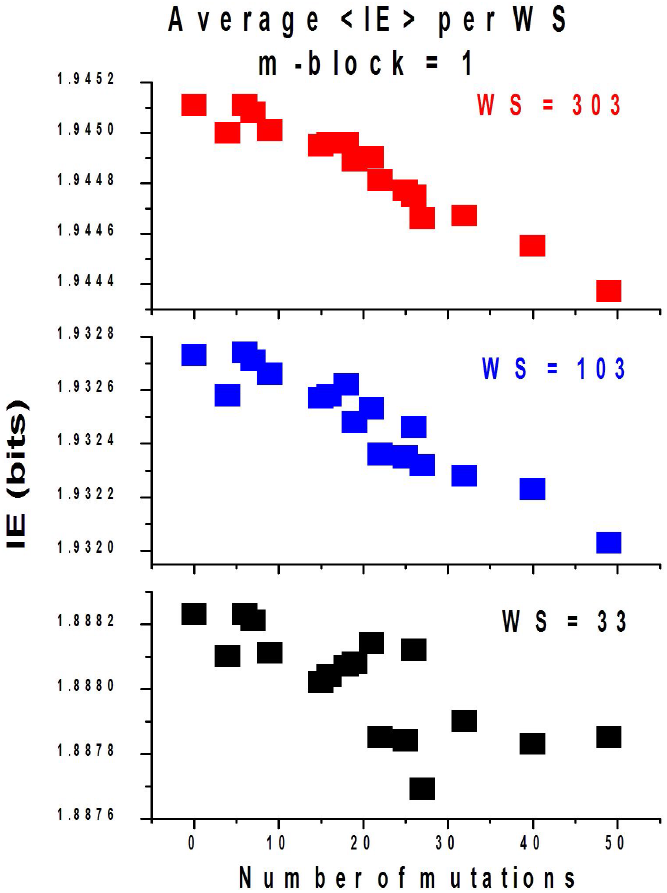
Average IE values per WS for each variant of SARS-CoV-2 studied, plotted as a function of the number of mutations.

**Figure 3.**
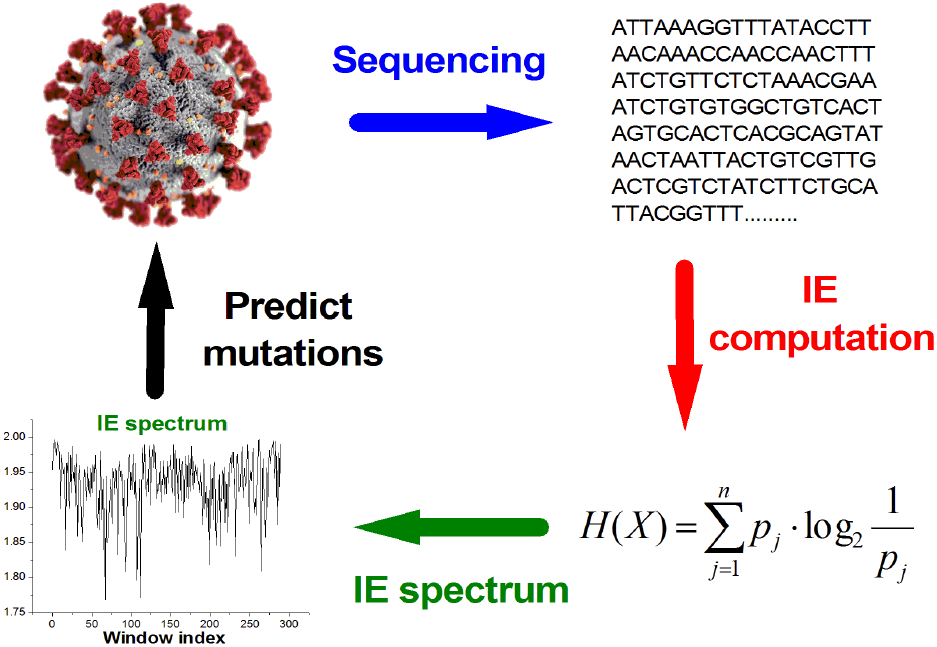
Diagrammatic representation of the methodology to produce the IE spectrum in order to facilitate studies of possible predictive algorithm of future genetic mutations.

different WS values, as a function of the number of mutations. The data indicates again that the average IE per window within each genome decreases with increasing the number of mutations. The larger the WS values, the better the correlation.

This confirms that splitting the genome into windows retains the correlation to the IE observed in figure 1, and the proposed governing law of genetic mutation is applicable to individual windows. In turn, this allows us to establish a link between the IE values and the index locations of the mutations, which could facilitate further studies to determine a full predictive algorithm of genetic mutations, as diagrammatically shown in figure 3.

To exemplify this technique, we computed the IE spectrum of the SARS-CoV-2 reference genome (MN908947) and the spectrum of the first mutated variant (LC542809) using WS = SS = 103. We performed a direct comparison of the two genomes and four SNP mutations were identified in the LC542809 sequence (see table 1), as following: G11082T, C14407T, C26110T and C29634T. The first character is the mutating nucleotide in the reference genome, the number is its location in the sequence and the second character is what it mutated into in the new sequence.

Figure 4 shows the obtained IE spectra of the two sequences and their IE ratio (IER), i.e. IER = (IE mutated) / (IE reference). Although the two spectra appear identical, their ratio spectrum obtained by dividing the mutated spectrum to the reference spectrum can identify consistent changes between the two genomes.

**Figure 4.**
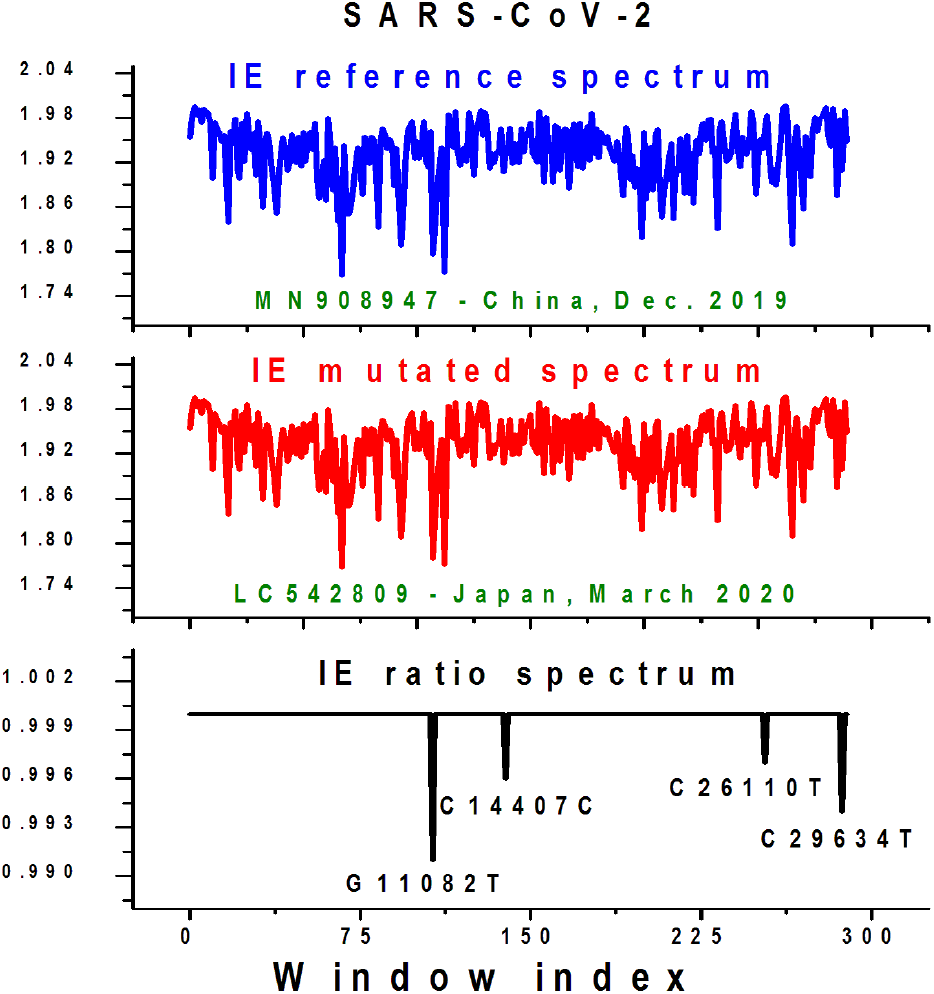
The IE spectra of the reference SARS-CoV-2 genome (top image), the mutated sequence (middle image) and their IE ratio (IER) spectrum (bottom image) obtained by dividing the two IE spectra. The four known mutations are captured in the IER spectrum and clearly marked. The IE of the corresponding window decreased in each case. Data obtained for WS = SS = 103.

Any value not equal to one in IER spectrum indicates the presence of at least one mutation inside the corresponding window of the spectrum, resulting in a change in the information encoded at that position. The IE ratio spectrum (figure 4, bottom image) successfully captured all four mutations and their corresponding locations within the genome’s IE spectrum, confirming the validity of this technique. Most importantly is the fact that the governing law of genetic mutations appears valid when applied to the individual windows where mutations have been detected, because each mutation resulted in a decrease of the IE value of the corresponding window, i.e. IER < 1, see figure 4.

Table 2 shows the extracted string of nucleotides corresponding to each window that suffered a mutation in the reference spectrum. The nucleotide that mutated is highlighted in red. We also show the corresponding IE value of each window before and after the mutation took place, indicating that mutations take place in a way that reduces the IE value within the selected window. Although in this example all four mutations resulted in a decrease of the IE values of their containing windows, it is important to mention that not all mutations display this behavior. Indeed, the majority of the mutations occur so that the IE value of their containing window decreases, but in some cases only 70-80% of the mutations obey this information entopic rule, especially for genomes that suffered a large number of mutations. When applied to the entire genomes, the observed entropic law is always fully applicable.

**Table 2.**
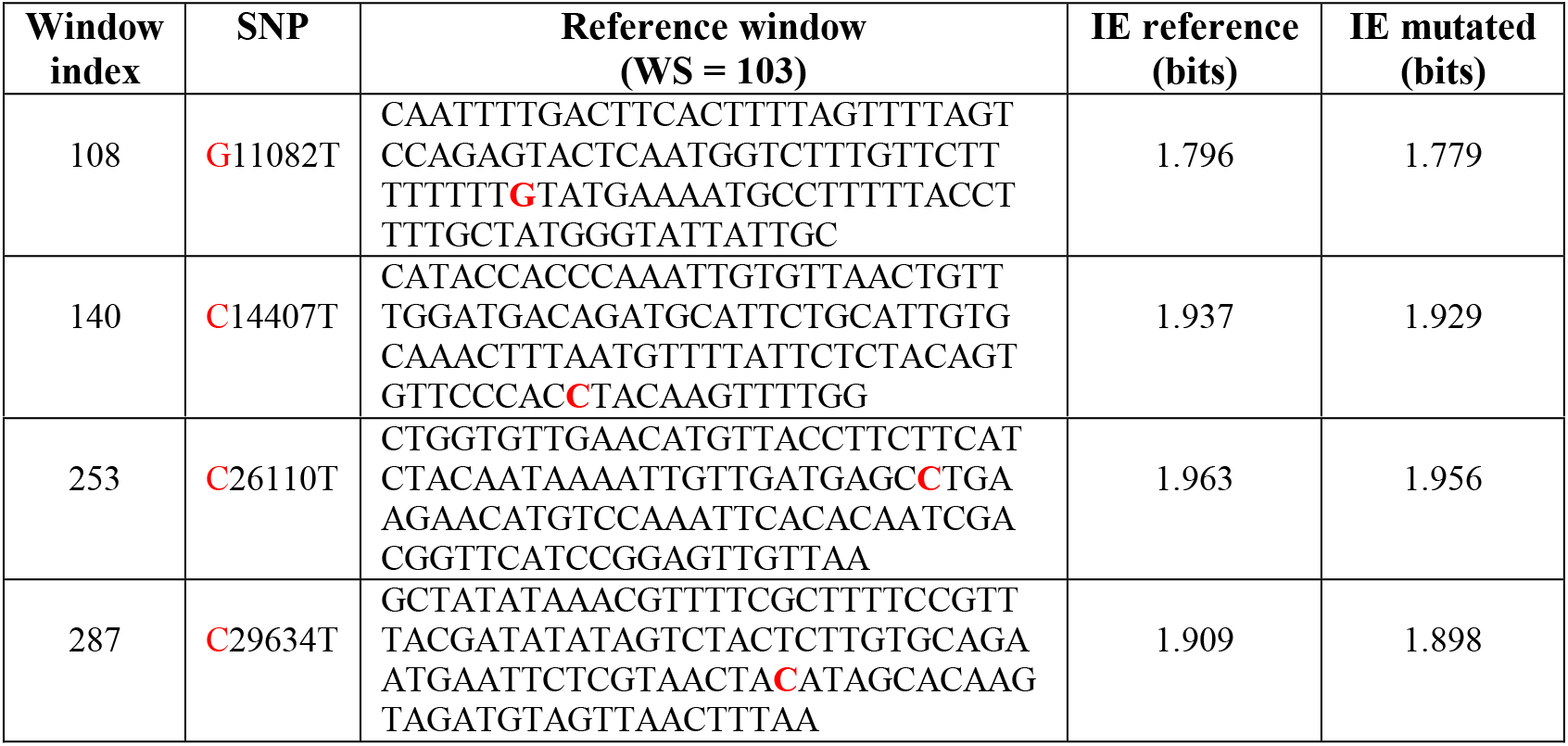
Details of the mutations that occurred in SARS-CoV-2 reference genome (MN908947) to result in the first mutated variant (LC542809), including the IE values of their corresponding windows where the mutations took place.

## Discussion

In conclusion we report here the observation of an entropic governing law of genetic mutations. The law states that genetic mutations are not mere random events, but they occur in a way that the information entropy (IE) of the whole genome reduces in value. In order to expand the applicability of this observation to the development of possible predictive algorithms of genetic mutations, we proposed a methodology that involves the conversion of the genetic sequences into IE numerical spectra by splitting them into segments of nucleotides called windows. The proposed entropic governing law of genetic mutations appears to be valid not only when applied to the full genome (figure 1), but also when applied to the average IE per window (figure 2), and most importantly when examining windows that contain specific mutations (figure 4 and table 2). This opens up the possibility of implementing this technique to obtain a full deterministic approach to genetic mutations. By carefully studying the IE spectrum of a genome and its variants, combined with the observed law of genetic mutations and machine learning algorithms, one should be able to establish a deterministic algorithm of predicting future mutations before they occur. This article focuses on mutations in the RNA of SARS-CoV-2 viruses, but we should keep in mind that DNA is subject to essentially the same entropic mutation forces and we expect the observations reported here to be valid in all biological systems.

## Methods

All the RNA sequences used in this study are freely available from the NCBI virus database (https://www.ncbi.nlm.nih.gov/labs/virus/vssi/#/). Each sequence has a reference download link shown in Table 1 of the manuscript. When extracting SARS-CoV-2 genetic sequences, only complete genomes with 29,903 characters should be downloaded, to match the size of the reference genome [7]. In order to use the GENIES code, FASTA (text) genome files should be formatted in a specific way that they contain no other characters except a long string of A, T, C and G letters. For this specific study, a modified version of GENIES has been used, in order to facilitate running the program with m-block size of one. This new program (GENIES_Jan_2022.vi) is also freely available to download from the same repository (https://sourceforge.net/projects/information-entropy-spectrum/), but there is no installer available and it requires LabView to run it. Currently, the main installer only allows minimum m-block size of two. The m-block is a group of characters (nucleotides) that form a new unique character within a given set. The choice of m-block size is very important as the maximum IE value per event is linked to the m-block size by the relation H(Y) = log_2_4^m-block^= 2 ×m-block. For example, if m-block size is one, then we can have only 4^1^= 4 possible occurrences (A, T, C and G) within a given genome or segment of genome and a maximum of H(Y) = 2 bits encoded per event. If we use m-block size two, then the number of unique possible events become 4^2^= 16 (AA, AT, AC, AG, TA, TT, TC, TG, CA, CT, CC, CG, GA, GT, GC, GG), so each event can encode maximum H(Y) = 4 bits and the number of unique events within a genome is given by the formula 4^m-block^. The current study is limited to single nucleotides, so m-block =1. In order to extract the IE per genome, one needs to use GENIES_Jan_2022 with the following input parameters: m-block =1, m-block step size = 1. The value of the window size is a variable and data is sensitive to this choice, so more research is required to understand the effect of the WS, but to reproduce the data shown here one should select WS = SS. In this article we used WS= 33, 103 and 303 to extract the average IE value per window and WS = 103 to produce the IE spectra and the IER spectrum for detection of genetic mutations. The IE values per whole genome have been obtained by selecting the WS = genome size = 29,903. Unless one needs to detect mutations, to extract these parameters, run GENIES by uploading the same sequence twice. If IER spectrum and mutations analysis are required, one should run the program following the on screen instructions. Detailed instructions on running the code and the procedure of computing the IE spectra are described in the GENIES user manual [9].

## Acknowledgments

MV acknowledges the financial support received for this research from the School of Mathematics and Physics, University of Portsmouth.

## Data Availability

The numerical data associated with this work is available within this manuscript. The RNA sequences used in this study are freely available from references 7 and 10-25. The code is freely available to download [8].

## Code Availability

The codes used in this study are freely available at: https://sourceforge.net/projects/information-entropy-spectrum/. Reference [9] also contains the user manual of this programme.

